# A resource for extracellular vesicles from activated CD4 T cells that relay pro-inflammatory signals

**DOI:** 10.1101/2024.10.15.618584

**Authors:** Ashwin K. Jainarayanan, Ranjeet Singh Mahla, Jesusa Capera, Mirudula Elanchezhian, Dhanu Gupta, Nithishwer Mouroug Anand, Tom Thomas, Daniel Beckers, Alexander Leithner, Sooraj R Achar, Salvatore Valvo, Mariana Conceição, Elke Kurz, Abhinandan Devaprasad, Sakina Amin, Georgina Berridge, Svenja Hester, Roman Fischer, Pablo F. Céspedes, Lynn B. Dustin, Matthew J. A. Wood, Michael L. Dustin

## Abstract

CD4 T helper cells (TH cells) play a vital role in coordinating and amplifying the immune response to specific pathogens. They constitutively produce different kinds of extracellular vesicles (EVs), which mediate cell-to cell communication and play diverse roles in immune regulation and inflammatory processes. Here we provide a resource documenting the composition of activated TH cell EVs and demonstrating their ability to instigate pro-inflammatory response in antigen-presenting cells (APCs). EVs were characterized by lipidomics, proteomics, and NanoFCM. The activated TH cells derived EVs (act-EVs) were found to be enriched in TH cell-specific proteins, transmembrane and cytosolic EV marker proteins, and HLA proteins relative to resting CD4 T cells EVs (rest-EVs). The pro-inflammatory effect of act-EVs vs rest-EVs on donor matched APCs were characterized by chemokine and cytokine profiling and flow cytometry analysis. There was no discernible contrast in endotoxin levels between act-EVs and rest-EVs. Functional distinctions were seen to arise from variations in the content and composition of these EVs. Moreover, we validated our findings with an in-vivo investigation in mice, demonstrating the recruitment of monocytes, dendritic cells (DCs), neutrophils, and NK cells in the spleen, accompanied by the release of pro-inflammatory cytokines in the serum after administering act-EVs. In summary, this study sheds light on the role of TH cell released EVs in modulating the immune response during pro-inflammatory responses and this resource provides a foundation for development of novel therapeutics on EV based scaffolds.

## Introduction

CD4 T helper cells (TH cells) are central in mounting adaptive and humoral immune responses in infection and inflammation. Through cytokine release, they activate and polarize many immune cells to ensure sufficient antibody production, engulfment and killing of infected cells, and further immune cell recruitment to the site of infection. Whilst cytokines are receptor-specific in their action, immune cells also release extracellular vesicles (EVs) which, whilst arguably less specific, are proving no less important in mediating an immune response (Dong, 2021). EVs are nanoscale vesicles which deliver multiple cargos to immune cells that can include DNA, RNA (Hirsch et al., 2023), lipids, glycans and proteins in compositions that overall, concisely reflect that of the parent cell (Kalluri, 2024). In this manner, EVs could be considered as waste material, yet in an immune context this ‘waste’ is recycled to fulfil an essential role in mediating cell-cell communication (Buzas, 2023; Ginini et al., 2022; Kalluri, 2024).

TH cells regulate the release of distinct exosomes, a subtype of EVs, under different activation contexts (van der Vlist et al., 2012). For instance, Cai *et a*l., showed that EVs from activated TH cells can induce the production of pro-inflammatory cytokines in myofibroblast to amplify the inflammatory response (Cai et al., 2020). Other studies suggest TH cell derived EVs may have anti-inflammatory effects. Upon activation, TH cell derived EVs regulate the immune response by modulating cytokine production in T cells, as well as monocytes and endothelial cells (Hezel et al., 2017; Scanu et al., 2008). In 2019, Torregrosa Paredes and colleagues found that EVs from regulatory TH cells can suppress the activation and proliferation of other immune cells, potentially reducing the severity of inflammation (Torregrosa Paredes et al., 2012). Overall, the role of TH cell derived EVs in an inflammatory environment is complex and likely depends on a variety of factors, including the physiological state of the TH cell producing the EVs, contents of the EVs, and the target cells that receive the EVs. Further research is needed to fully understand the role of TH cell derived EVs in mediating pro-inflammatory responses and their potential as therapeutic agents.

Interestingly, TH cell derived exosomes are reported to contain cell adhesion molecules and kinases that can facilitate the binding of EVs to target cells and initiate downstream signaling pathways, respectively (Hong et al., 2017). Such targeting, combined with the capacity to carry cytokines, chemokines, and other molecules in their proteinaceous coronas that enhance the immune response and promote cell survival, center EVs as potent immunomodulatory agents to enhance immune responses against autoimmunity, cancer, and infectious diseases (Kalluri, 2024). With their anticipated therapeutic potential, it is essential to understand the biological compositions of TH cell derived EVs.

In this study, we characterized the EVs derived from activated TH cells (act-EVs) and resting TH cells (rest-EVs) through proteomic and lipidomic analyses. Size and structural features of act-EVs and rest-EVs were examined using nanoparticle tracking analysis (NTA), NanoFCM, and direct STochastic Optical Reconstruction Microscopy (dSTORM). The functionality of act-EVs and rest-EVs in their respective donor matched recipient cells (APCs such as B-cells and monocytes) were analysed using cytokine (cytokine and chemokine array), and activation marker (flow cytometry) profiles. Additionally, we corroborated our findings through an in-vivo study which revelated the recruitment of monocytes, DCs, neutrophils, and NK cells (in spleen) along with the release of pro-inflammatory cytokines (in serum) upon act-EVs administration. This study provides extensive insights into the biology of EVs derived from TH cells of physiological states including their characterization, functionality and potential to modulate immune regulation.

## Methods and Materials

### Mice

FVB mice were housed under pathogen-free conditions in the Institute of Developmental and Regenerative Medicine or Kennedy Instiute of Rheumatology Animal Facility in accordance with local and UK Home Office regulations. The mice were housed in individually ventilated cages (IVC) with corn cob bedding. 12 hours of light/dark cycle with half an hour of dim light period in place. Appropriate environmental enrichment was provided: Enviro-dri, and housing tunnel, cage balcony and chew blocks. The temperature was maintained at 21 degrees +/-2 and the humidity 55% +/- 10. The protocols were reviewed by local Veterinary Surgeon (Vet) and Named Animal Care and Welfare Officer (NACWO) before being reviewed and approved by Animal Welfare and Ethical Review Body (AWERB). All procedures were conducted in accordance with the UK Scientific Procedures Act of 1986 and overseen by University of Oxford Department of Biomedical Services, Clinical Medicine Old Road Campus.

### Splenocyte preparation

Spleens removed aseptically from CO_2_ asphyxiated FVB mice was placed in sterile 100 mm dish and washed with PBS, spleen was minced into fine pieces, and passed through 70 µm cell strainer into a 50 mL tube using sterile plunger, the RBCs were lysed by RBC lysis buffer, lysis stopped by adding RPMI. Cells were washed for 5 mins at 500g and cryopreserved in 90% FBS 10% DMSO until further use.

### Human Peripheral Blood cells

The conducted research complies with the relevant ethical regulations at the University of Oxford. Different T cell populations were isolated from leukapheresis reduction system (LRS) chambers from de-identified, non-clinical and consenting healthy donors. The non-clinical issue division of the National Health Service and the Inter-Divisional Research Ethics Committee of the University of Oxford approved the use of LRS chambers (REC 11/H0711/7 and R51997/RE001).

### Isolation and purification of conventional TH cells

The untouched TH cells were isolated from LRS peripheral blood mononuclear cells (PBMCs) using RosetteSep™ Human CD4+ T Cell Enrichment Cocktail (StemCell Technologies, #15022, #15062). The negatively selected TH cells were activated with anti-CD3/CD28 Antibody-coupled Magnetic Beads (ThermoFisher # MBS-C001-10mg) at a cell-to-Dynabeads ratio of 1:1 and 100 U/mL of IL-2. At 48 hrs after activation of cells, the CD3/CD28 Dynabeads were removed from the cells and cells were maintained in complete RPMI-1640 (RPMI 1640 medium supplemented with 10% heat-inactivated fetal bovine serum, 100μM non-essential amino acids, 10mM HEPES, 2mM L-glutamine, 1mM sodium pyruvate, 100U/mL of penicillin and 100μg/mL of streptomycin) with 100 U/mL of IL-2. Following this, RPMI-1640 was replaced with OptiMEM (Gibco, OptiMEM) and the cells were maintained for 48 hrs before isolating the constitutively released EVs, as mentioned in (Jainarayanan et al., 2023).

### Size-exclusion liquid chromatography (SELC)

The EVs were isolated from the OptiMEM media using SELC method (EV isolation protocol standardisation discussed in (Jainarayanan et al., 2023)). The cells along with culture supernatant were transferred to 15 ml Falcon tubes and cells were pelleted at 300g for 10 mins. Cell and debris-cleared supernatants were transferred to fresh 15 ml Falcon tubes and centrifuged at 4,000g for 10 mins, pelleting cell debris. Next, the supernatant was concentrated using 100 kDa molecular weight cut-off Amicon centrifugal filter units (Merck Millipore) to 2 ml volume, loaded to Sepharose 4 Fast Flow affinity resin column (10 mm x 300 mm; GE Healthcare) mounted to ÄKTA pure system (GE Healthcare) and eluted at 0.5 ml/min using PBS as the eluent. The elution chromatogram was recorded at 280 nm and each elution volume was set to 2 ml. The EV-containing elution fractions were pooled and concentrated using 10 kDa MW cut-off Amicon centrifugal filter units (Merck Millipore # UFC5010).

### Western blotting

Whole cell lysates were prepared by lysing TH cell pellets in RIPA buffer (Thermo Fisher Scientific, #89901) containing Protease/Phosphatase inhibitor (PI) cocktail (CST; #5872) to a final concentration of 2 x 10^7^ cells/mL. The lysed cells were centrifuged at 10,000g for 10 mins at 4° C to remove cellular debris, and the protein supernatants were collected. The protein concentration was determined by BCA, protein samples were denatured in Laemmli buffer at 95° C for 10 mins and subsequently loaded on a SDS PAGE gel. In case of particles, isolated vesicles were subjected to RIPA and PI treatment, the protein concentrations were measured using micro-BCA and equal amounts of proteins across EV samples were loaded. Samples were resolved using 4-15% Mini-PROTEAN SDS-PAGE gels (Bio-Rad; #4561084), and transferred to 0.45 μm nitrocellulose membranes (Bio-Rad, #1620115), blocked with 5% BSA and incubated with the following primary antibodies: mouse anti-ALIX clone 3A9 (CST, #2171), rabbit anti-APOA2 clone EP2912 (#ab119990), rabbit anti-CD4 clone EPR6855 (#ab133616), rabbit anti-CD40 Ligand clone D5J9Y (CST, #15094), rabbit anti-ALB clone EPR20195 (#ab207327), rabbit anti-CD69 clone EPR21814 (#ab233396), rabbit anti-calnexin clone C5C9 (CST, #2679), rabbit anti-CD3 zeta clone EP286Y (#ab40804), mouse anti-CD81 clone M38 (Invitrogen, #10630D), rabbit anti-HIST1H1.0 clone EPR6537 (#ab125027), mouse anti-LMNA clone 133A2 (#ab8980), rabbit anti-TSG101 clone EPR7130B (#ab125011). Following incubation with primary antibodies, membranes were washed thrice with 1X TBST wash buffer and incubated with IRDye 800CW donkey anti-rabbit IgG (H+L; LI-COR, #925-32213) and IRDye® 680RD donkey anti-mouse IgG (H+L; LI-COR, #926-68072) secondary antibodies. Primary and secondary antibody dilutions were used according to the manufacturer’s instructions. Membrane blocking with 5% BSA, or incubation with primary or secondary antibodies were performed either overnight at 4°C or 1 hr at room temperature. After incubation with secondary antibodies, membranes were washed four times with 1X TBST and image acquisition was performed using the Odyssey® CLx Near-Infrared detection system operated with Image Studio™ Lite quantification software (LICOR, Lincoln, NE).

### NanoParticle Tracking Analysis (NTA)

The isolated particles were diluted in PBS to reach a concentration of 5E+8 to 1E+9 particles per ml. The instrument used for NTA was Nanosight NS500 (Malvern Instruments Ltd), together with NTA 3.2 software, set to light scattering mode. Videos for NTA were captured for 60 s, three sequential replicates per sample were obtained, and the three recordings were processed and averaged to determine the mean size and concentration of the particles. For the tracking analysis, the TrackMate plugin from ImageJ was used (Tinevez et al., 2017). A mask video was created from the raw videos to detect the spots in each frame. To create the mask video, each frame was processed with background subtraction using the rolling-ball method, followed by filtering using Gaussian Blur and Median filters. Next, a threshold for each frame was set using the Bernsen local thresholding method and converted into a binary image. Detected spots in the mask video were filtered by area (>40 nm^2^) and circularity (>0.5). Frame-to-frame spot linking was performed using a Linear Assignment Problem tracker with a maximum linking distance of 20 nm, a gap-closing maximum distance of 30 nm and a gap-closing maximum frame gap of 4. Only tracks with a minimum length of 30 frames were considered.

### Transmission electron microscopy (TEM)

For TEM, the thinly spread vesicle preparations were negatively stained as previously described (Harris, 2007). In brief, copper grids (300 mesh) with carbon support film coating of 3mm were plasma treated for 20 s using a Leica EM ACE200 Vacuum Coater. Next, 10 µL of the isolated vesicle populations were deposited on the grids and incubated at room temperature for 5 mins. The excess sample was removed with a Whatman N°1 paper, and the samples were stained at room temperature with 2% of uranyl acetate 10 seconds. After removing the excess uranyl acetate, the samples were dried for 10 mins and imaged using a Tecnai 12 TEM at 120 kV with a Gatan OneView CMOS camera. The imaging was carried out at a final magnification of 29,000x for isolated vesicle populations.

### Nano Flow Cytometry

The Nano flow cytometry analysis was performed using the Flow NanoAnalyzer (NanoFCM Co., LTD) according to the manufacturer’s instructions. Silica Nanospheres Cocktail (S16MExo, NanoFCM) was employed as the size standard to construct a calibration curve to convert side scatter intensities to particle size. NanoFCM 200nm PS QC beads were used as a concentration standard for calculating particle concentration in EV samples. Antibodies were allowed to bind for 30 mins on ice. Unbound antibodies were removed by centrifugation for 1h at 100,000g at 4 °C. Labelled vesicles were then resuspended in 50 µL of PBS. The laser used was a 488nm laser at 25/40 mW, 10% ss decay. Detectors were equipped with 525/40 (AF488) and 580/40 (PE) bandpass filters. Buffer alone (PBS), unstained samples, samples stained with isotype controls and auto-thresholding were used to define sample and label specific signals.

### Total Internal Reflection Fluorescent Microscopy (TIRFM) and dSTORM imaging

TIRFM was performed on an Olympus IX83 inverted microscope equipped with a TIRF module. The instrument was equipped with an Olympus UApON 150 x 1.45 NA oil-immersion objective and Photomertrics Evolve delta EMCCD camera. Image analysis and visualization were performed using ImageJ (National Institutes of Health) (Schneider et al., 2012). For TIRFM, the EVs were labelled by membrane dyes DiD/DiI (ThermoFisher Scientific; #V22887/#V22888) and Wheat Germ Agglutinin (WGA) conjugated with AlexaFluor488 (ThermoFisher Scientific; #W11261) and captured onto poly-L lysine coated glass (Balint et al., 2020). For dSTORM imaging EV samples were mounted with a reducing buffer system. A total of 10,000 images were captured on a Nanoimager (Oxford Nanoimaging) with a 100x oil-objective lens, and captured images were analyzed with Nanoimager Software v1.4.8740 (Oxford Nanoimaging).

### Multicolour Flow Cytometry (FCM)

The staining was performed in 5% BSA in PBS pH 7.4 (0.22 µm-filtered) at +4°C for at least 30 mins. Further, cells were washed thrice and then acquired using an LSRFortessa X-20 flow cytometer with a High Throughput Sampler (HTS). Fluorescence spectral overlap compensation was then performed using single colour-labelled cells and unlabelled cells. To calculate compensation matrixes for markers with low surface expression levels, unstained and single colour stained UltraComp eBeads were used. The antibodies used for staining human APCs were anti-CD54-BV711 clone HA58 (BD Bioscience, #564078), anti-HLA-DR-AF488 clone L243 (Biolegend, #307619), anti-CD40-PerCP/Cy5.5 clone HB14 (Biolegend, #313013), anti-CD83-PE/Cy7 clone HB15e (Biolegend, #305325).

For mouse splenocyte staining, the cells were resuspended in 100 µL of Live/Dead eFluor780 (eBioscience, #65086514) diluted 1:1000 in PBS for 10 min at room temperature. Cells were washed with FACS buffer and incubated for 20 min with the staining cocktail consisting of antibodies. The antibodies used in the staining cocktail were anti-NK1.1-BV421 clone PK136 (Biolegend, #108731), anti-CD14-BV605 clone Sa14-2 (Biolegend, #123335), anti-Ly-6C-BV711 clone HK1.4 (Biolegend, #128037), anti-CD11b-FITC clone M1/70 (Biolegend, #101205), anti-Ly-6G-PE clone 1A8 (Biolegend, #127607), anti-CD11c-PE/Cy5 clone N418 (Biolegend, #117316), anti-CD49b-PE/Cy7 clone DX5 (Biolegend, #108921), anti-CD4-PerCP/Cy5.5 clone GK1.5 (Biolegend, #100433), anti-CD8a-PE/Dz594 clone 53-6.7 (Biolegend, #100761), anti-TCRb-AF647 clone H57-597 (Biolegend, #109217), anti-B220-AF700 clone RA3-6B2 (Biolegend, #103231). After incubation, cells were washed twice with FACS buffer and resuspended in R10 for sort. The gating strategy is shown in Figure 5B.

### Fluorescence-activated cell sorting (FACS)

For staining human PBMCs, the cells were resuspended in 100 µL of Live/Dead eFluor780 (eBioscience, #65086514) diluted 1:1000 in PBS for 10 min at room temperature. Cells were washed with FACS buffer and incubated for 20 min with the staining cocktail consisting of antibodies. The antibodies used in the staining cocktail are anti-CD19-AF700 clone SJ25C1 (Biolegend, #363033), anti-TCRα/β-BV421 clone IP26 (Biolegend, #306721), anti-CD14-FITC clone HCD14 (Biolegend, #325603), anti-CD66b-APC-A clone G10F5 (Biolegend, #305117). After incubation, cells were washed twice with FACS buffer and resuspended in R10 for sort. The gating strategy is shown in Supplementary Figure 4A. All sorts were performed using a BD FACSAria™ III cell sorter and data were analyzed using Flowjo v10.7.1.

### Human Cytokine Arrays and Image Analysis

Human cytokine profiling was performed using a Proteome Profiler Human XL Cytokine Array Kit (R&D Systems, #ARY022B), which detects 102 human soluble cytokines. Human XL Cytokine Arrays were incubated overnight at 4 °C with 1.5 mL of the cell culture supernatant, and the procedure was performed according to the manufacturer’s instructions. Following incubation with a detection antibody cocktail, antibody conjugation, and recommended washes, the immunoblots on the membrane were developed with Chemiluminescent Substrate Reagent Kit (Invitrogen, # WP20005) and were exposed to X-ray film. Cytokine array TIF file images were analyzed using ImageJ. After uploading images into the software, a grid file was manually superimposed to assign spot gene identities. In the next step, spot mean intensities were measured. For background correction, median background intensities were subtracted from mean signal intensities. To further calculate the differential expression, the mean signal intensity of the cytokine/chemokine spots was divided by the mean signal intensity of the reference spots.

### scRNA sequencing

The raw scRNA-seq data was downloaded from Gene Expression omnibus (GEO) under accession number GSE126030 (Szabo et al., 2019). Only 4 samples corresponding to PBMCs from this accession code were used for the analysis. This includes the GEO accession: GSM3589418, GSM3589419, GSM3589420 and GSM3589421. The T cells from these PBMCs were activated using Human CD3/CD28 T Cell Activator (STEMCELL Technologies) and compared to the resting T cells cultured in media. The fastq files obtained from GEO were processed using the Cell Ranger (v3.1.0) counts pipeline. The sequencing reads were aligned to Human GRCh38-3.0.0 and the cell count matrices were obtained. These filtered genes–cell matrices were then read into the Seurat v4.0 pipeline implemented in R for downstream analysis. Although we used the same dataset, we re-clustered the data, and the cells were reannotated to examine the cell types of interest. We followed the standard Seurat pre-processing workflow with QC-based cell filtration, data normalization and scaling. Low-quality cells with less than 200 unique feature counts and cells with a mitochondrial count greater than 5% were excluded. We ended up with roughly 6000 cells in the processed dataset. We further identified major TH cell types with highly variable genes and performed Jackstraw analysis to isolate significant principal components. The UMAP embeddings were then computed using the isolated PCs and known marker genes were used to characterize cell types.

### Proteomics

The EV samples were reduced with 5 μL of 10 mM tris(2-carboxyethyl) phosphine and alkylated with 50 mM of iodoacetamide for 30 min each, acidified with 12% phosphoric acid 10:1 vol:vol, and then transferred to S-trap columns. Samples were precipitated using 1:8 vol:vol dilution of each sample in 90% methanol in 100 mM Triethylamonium bicarbonate and digested with trypsin (Promega, #V1115) overnight at 37° C. The samples were then run on a LC-MS system comprised of an Evosep One and Bruker timsTOF Pro. For proteomic analyses, the raw files were searched against the reviewed Uniprot Homo sapiens database (retrieved 2,01,80,131) using MaxQuant v1.6.10.43. The LFQ intensities were normalized using particle concentration from NTA and further average-based normalization was carried out to obtain the enriched set of proteins in each EV sample. In addition, for human plasma proteomics average based enrichment was carried out (Deutsch et al., 2021).

### Lipidomics

The extraction process began by taking approximately 250 µL of each sample. Then, 200 µL of cold methanol followed by 800 µL of MTBE was added to each sample. The sample was mixed on a horizontal mixer for about 6 mins. This was followed by adding 200 µL of H_2_O and centrifugation at 10,000g for 10 mins at 4°C. After centrifugation, the top layer was dried in a Speedvac for ∼2 hrs. The pellet was resuspended into 15 µL of lipid resuspension solution (ButOH:IPA:H2O (8:23:69)), and 1 µL was loaded to MS/MS for high mass accuracy analysis. The intensities were normalized using the particle concentration based on NTA for the lipidomic analysis. The lipidomics analysis was performed using Lipid Ontology (LION) tool.

### Endotoxin Assay

LPS levels were measured with Pierce Chromogenic Endotoxin Quantification Kit (ThermoFisher, #A39552) as per manufacturer’s instructions.

### In-vivo Studies

Wildtype FVB mice (n=3) were utilized in this study. Animals were euthanised using CO2 inhalation and their spleens were aseptically extracted to isolate TH cells employing MojoSort™ Mouse CD4+ T Cell Isolation Kit (Biolegend, #480033). Isolated T cells were cultured in complete RPMI-1640 medium and activated using CD3/CD28 Dynabeads to induce activation. EVs (act-EVs) were obtained from the culture media using the SELC method post 48 hrs. For control, 200 μL of freshly isolated mice plasma was diluted with 4.8 mL of PBS and was centrifuged for 15 mins at 2000g to remove cells and platelets, and supernatants were aliquoted into fresh tubes and EVs (plasma-EVs) were isolated using SELC. Approximately 0.8 x 1011 of EVs were administered into the FVB mice (n=3) via tail vein injection. Blood samples were collected at 4 hours post-EV administration to assess cytokine profiles in the serum. The levels of various cytokines were quantified using LEGENDPLEX (BioLegend, #740446) to analyze the impact of EV administration on cytokine profiles. Further FCM was performed to quantify diverse immune cell populations retrieved from the spleen of mice 4 hours after the administration of EVs.

### Statistical analysis

Statistical analysis was conducted utilizing GraphPad Prism software v10.0.3. To assess statistical significance, various methods were employed, including normality tests like Kolmogorov-Smirnov or Shapiro-Wilk tests, followed by appropriate tests such as Mann-Whitney test, unpaired t-test, nested T-test or Two-way ANOVA Šídák method.

## Results

### Upregulation of EV biogenesis with increased proportion of TCR+ EVs in TH cells upon anti-CD3/CD28 stimulation

The isolation of act-EVs and rest-EVs was achieved using SELC, as depicted in Figure 1A. The particle sizes in resting and activated TH cells were characterized through NTA, revealing that particles obtained from activated TH cells exhibited a higher concentration (peak concentration of 8 x 10^8 particles per ml) compared to those isolated from resting TH cells (peak concentration of 4.5 x 10^7 particles per ml), as illustrated in Figure 1B and 1C. This observation was qualitatively validated using TEM as shown in Figure 1D. This was in coherence with scRNA-seq analysis, which demonstrated the upregulation of genes associated with both T cell activation and EV biogenesis in activated TH cells compared to resting TH cells (Supplementary Figure 1C and 1F). Furthermore, genes involved in calcium metabolism and EV release were enriched in memory TH cells, while those involved in TCR signalling and kinase mediated signalling were enriched in effector TH cells (Supplementary Figure 1D and 1G). KEGG pathway analysis for the top 50 differentially expressed genes in activated TH cells revealed pathways such as glycolysis, antigen processing and presentation, and oxidative metabolism (Supplementary Figure 1E).

**Figure 1:**
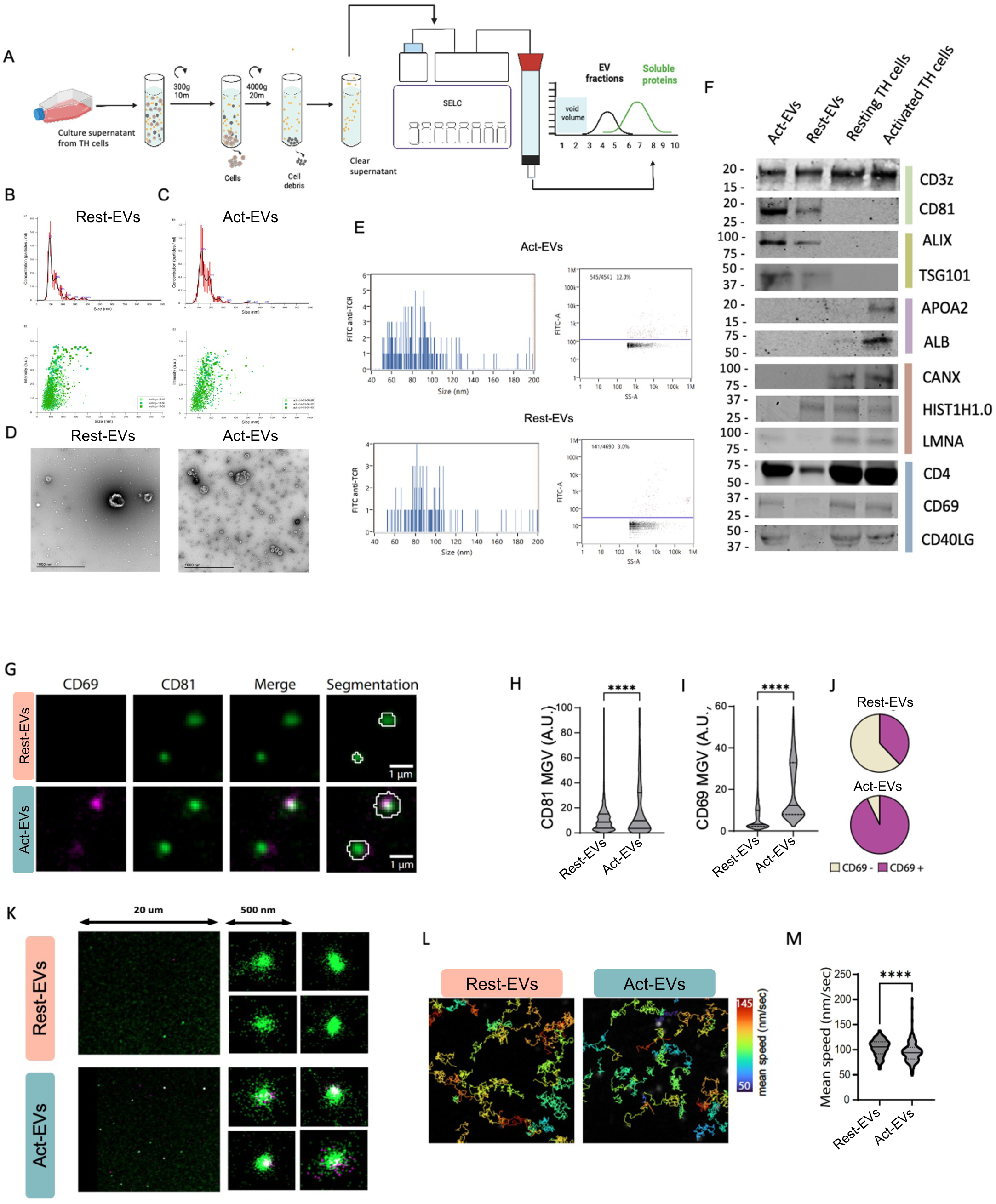
(A) Schematic showing the EV isolation methodology. (B-C) NTA analysis showing the concentration and size distribution rest-EVs (B) and act-EVs (C). (D) Transmitted electron microscopy images of eluates obtained using SELC EV isolation technique showcasing EVs and soluble proteins isolated from activated (Right panel) and resting TH cells (Left panel). (E) NanoFCM analysis showing the scatterplot for TCR positive EVs marked in red above the blue line and the others below. Frequency of TCR positive events recorded on the stained rest-EVs and act-EVs. (F) Western blot image for the lysed isolated rest-EVs and act-EVs with their cell lysate labeled for transmembrane proteins (light green), cytosolic proteins (olive green), proteins that are major components of non-EV co-isolated structures (purple), transmembrane/ lipid-bound/ soluble proteins associated with other intracellular compartments (brown), and TH cell related proteins (blue). (G) Purified rest-EVs and act-EVs were immunostained for CD69 (magenta) and CD81 (green), and imaged with TIRFM. CD81 positive and/or CD69 positive particles were segmented and mean intensity grey values (MGV) for CD81 and CD69 were analysed within each particle. (H-I) Violin plots showing the distribution of the mean intensity of CD81 (H) and CD69 (I) for the particles detected in the TIRFM images of rest-EVs and act-EVs. Data shows the frequency distribution of >1,500 particles. Lines show the median and quartiles. (J) The percentage of CD69 positive (CD69+) and negative (CD69-) particles was calculated from all particles detected in rest-EVs and act-EVs. (K) Representative two-colour (CD81 in Green and CD69 in Meganta) dSTORM images of rest-EVs and act-EVs. (L) Representative images showing the tracks from rest-EVs and act-EVs in solution. Code colour shows the mean speed (nm/sec) of the particles across the tracks. (M) Violin plots showing the distribution of the mean speed (nm/sec) of the tracks from rest-EVs and act-EVs. Data shows the frequency distribution of >100 tracks. Lines show the median and quartiles. Mann Whitney test was performed, ****pv<0.001.

NanoFCM was employed to validate the findings, revealing a higher abundance of T cell receptor-positive (TCR+) act-EVs (12%) compared to rest-EVs (3%), as depicted in Figure 1E. The enrichment of transmembrane and cytosolic EV marker proteins, as well as TH effector-specific proteins, was observed in act-EVs compared to rest-EVs. In accordance with the Minimal Information for Studies of Extracellular Vesicles guidelines (Thery et al., 2018), most non-EV marker proteins (such as ALB, APOA2, HIST1H1.0, LMNA, and CANX) were relatively more depleted in both act-EVs and rest-EVs, as illustrated in Figure 1F.

### Distinct biochemical and physical properties of act-EVs

To delve deeper into the characterization of rest-EVs and act-EVs, samples were subjected to staining for CD69 (activation marker) and CD81 (EV marker), as depicted in Figure 1G. Notably, CD81 and CD69 exhibited significant enrichment in act-EVs, as illustrated in Figure 1H and 1I. The percentage of CD69+ EVs also demonstrated an increase upon stimulation, as shown in Figure 1J. This finding was further substantiated through dSTORM imaging of the samples, depicted in Figure 1K. Furthermore, the physical behaviour of the EVs in suspension was assessed using single-particle tracking analysis, as demonstrated in Figure 1L. Intriguingly, it was observed that act-EVs exhibited a slower movement compared to rest-EVs, as indicated in Figure 1M.

### Transmembrane proteins, signaling factors, adhesion molecules and kinases in activated TH cells are mirrored in the EVs profile

Furthermore, proteomic analysis (Supplementary File 1 & 2) was performed to unveil the enrichment profile of EVs, revealing that only 63 and 75 proteins were uniquely enriched in act-EVs or rest-EVs, respectively (Figure 2A). It was interesting to observe the abundance of plasma proteins in act-EVs (Supplementary Figure 2B), showing their ability to associate with plasma proteins (for example, IGHA1, IGHG1). The expression level of proteins in act-EVs surpassed that in rest-EVs (Figure 2B). Principal component analysis (PCA) demonstrated a distinct protein profile for act-EVs and rest-EVs (Figure 2C). Expectedly, proteins associated with T cell activation, such as IL2, CD40LG associated with pro-inflammatory response (Chen et al., 2006; Zirlik et al., 2007), T cell EV markers CD9 and CD81, effector proteins enriched in transynaptic vesicles like GRB2 and ADAM10 (Cespedes et al., 2022) were differentially upregulated in act-EVs. Interestingly, CD5, which negatively regulates TCR signalling from the onset of T cell signalling was also found to be increased in EVs upon TH cell activation (Perez-Villar et al., 1999) (Figure 2D). The expression profile revealed a higher abundance of tetraspanins, adhesion molecules, antigen-presenting molecules, transmembrane proteins, and cytoskeletal proteins in act-EVs compared to rest-EVs (Figure 2E) (Colombo et al., 2014; Jankovicova et al., 2020; Jurj et al., 2020; Willms et al., 2018).

**Figure 2:**
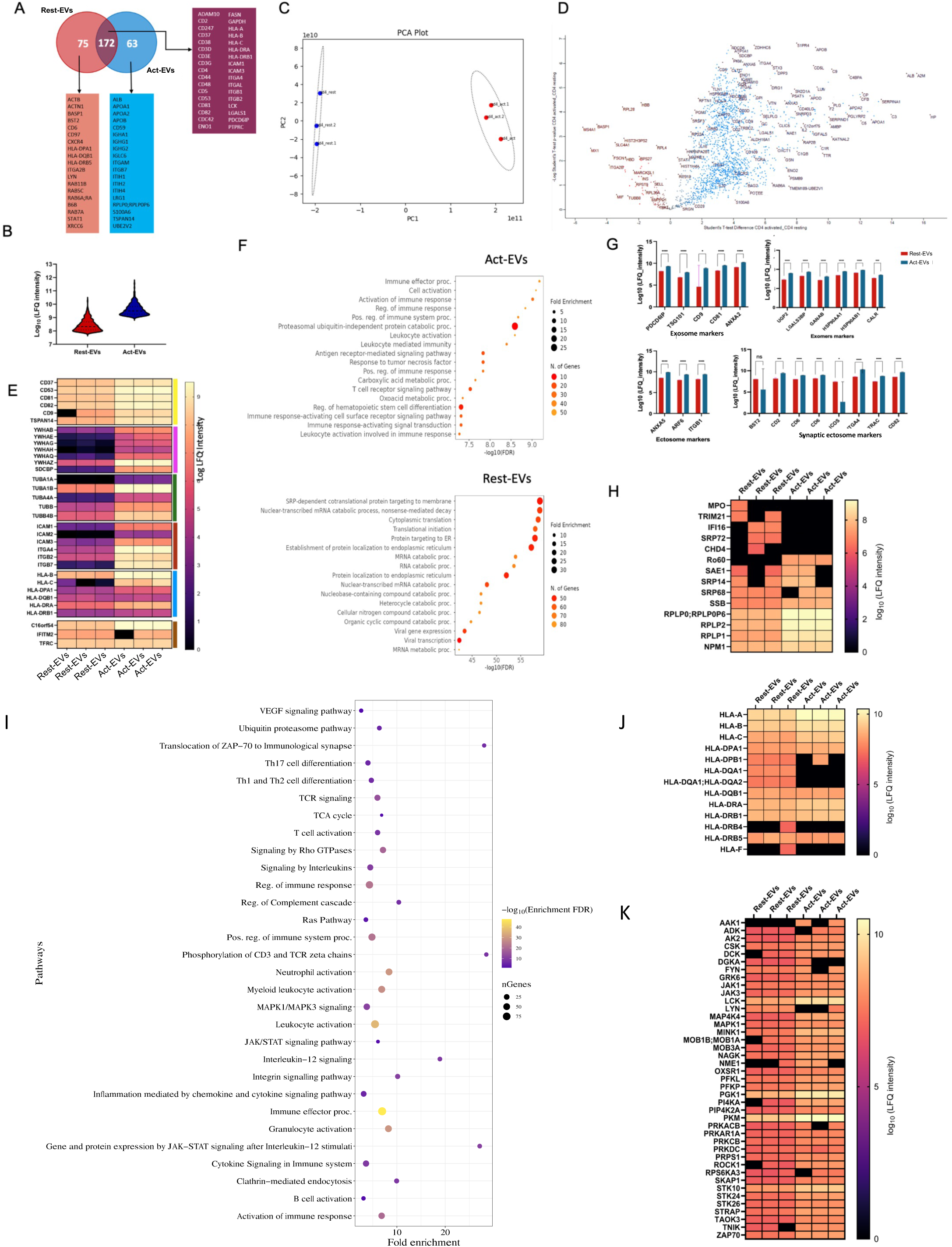
(A) Venn diagram depicting the numbers of shared and unique enriched proteins (some of the proteins are listed down) between the rest-EVs and act-EVs. (B) Violin plot showing the distribution of the protein contents in the rest-EVs and act-EVs. Lines show the median and quartiles. (C) PCA biplot of the samples showing variation in the protein profiles between the rest-EVs and act-EVs. (D) Volcano plot representing the differentially expressed proteins in rest-EVs (Red) and act-EVs (Blue). (E) Heatmap depicting the expression of different major protein classes in the rest-EVs and act-EVs. The major proteins of different protein classes, namely tetraspanins (yellow), signal transduction proteins (pink), cytoskeletal proteins (CPs) (green), adhesion molecules (AMs) (red), antigen-presenting molecules (APMs) (sky blue) and transmembrane proteins (TMPs) (brown). (F) Gene Ontology (GO) biological processes associated with proteins released by act-EVs (Top panel) and rest-EVs (Bottom panel). (G) Bar charts showing the expression of different EV class: exosome, exomere, ectosome and synaptic ectosome markers across the rest-EVs and act-EVs. Unpaired T-test was performed. (H) Expression of autoantigens in rest-EVs and act-EVs. (I) Reactome pathway analysis for differentially enriched genes from act-EVs. (J-K) Expression level of HLA proteins (J) and protein kinases (K) in rest-EVs and act-EVs. Data are representative of n =3 individual replicates and the LFQ intensities were log10 transformed. The significance level is *p≤0.05, **p≤0.01, ***p≤0.001 and ****p≤0.0001.

Signaling-related proteins, including the calcium-dependent kinase ZAP70, were predominantly enriched in act-EVs. Other enriched proteins encompassed SPN (CD43), essential for regulating T cell proliferation and migration (Mody et al., 2007), and FLNA, involved in the regulation of force transmission through integrins in T cells and T cell shear flow adhesion, crucial for T cell homing and trafficking into sites of inflammation (Savinko et al., 2018) (Supplementary Figure 3A). Following these results, an analysis of biological processes associated with proteins in act-EVs and rest-EVs (Figure 2F) revealed pathways related to leukocyte activation-related processes, antigen receptor-mediated signalling pathway and proteasomal ubiquitin-independent protein catabolic processes associated with act-EVs. On the other hand, translation-related pathways and protein localization to endoplasmic reticulum pathways were associated with rest-EVs.

Several markers associated with different types of EVs, such as exosome, exomere, ectosome, and synaptic ectosome, were significantly enriched in act-EVs (Figure 2G) (Jaiswal and Sedger, 2019; Kalra et al., 2016; Kim et al., 2017; Muralidharan-Chari et al., 2009; Saliba et al., 2019; Sheehan and D’Souza-Schorey, 2019; Zhang et al., 2018), suggesting that not only EV biogenesis increases but also different types of EVs are released upon activation of TH cells. Act-EVs also showed enrichment of RNA-binding proteins (Figure 2H). Reactome pathway analysis of differentially enriched proteins from act-EVs revealed pathways involved in ’TCR signaling,’ ’Translocation of ZAP70 to immunological synapse,’ ’Phosphorylation of CD3 and Zeta chains,’ ’B cell activation,’ ’Interleukin-12 signaling,’ and ’Regulation of the immune system’ (Figure 2I), indicating the potential role of EVs in cellular communication and downstream signaling pathways. The cellular component analysis showed that act-EVs (Supplementary Figure 3B) were predominantly from lysosome and complement membrane attack complex (C5-C9). In contrast, rest-EVs (Supplementary Figure 3C) were mostly ribosome-associated, senescence and telomerase-associated.

Most of the HLA proteins and protein kinases were enriched in act-EVs but not in rest-EVs. The enrichment in HLA proteins implies that the EVs are potentially capable of binding APCs. Protein kinases such as FYN and LYN, that act as molecular switches to direct immunoreceptors either towards homeostasis or inflammation (Mkaddem et al., 2017) are enriched in act-EVs and rest-EVs, respectively, indicating differential regulatory roles (Figure 2J and 2K). Apolipoproteins were enriched in EVs upon TH cell activation and some of these proteins are pro-inflammatory such as APOC3 and APOM, while some are known to regulate T cell activation such as APOA1, APOE, and APOA4 (Ruiz et al., 2017; Tenger and Zhou, 2003; Zewinger et al., 2020). CD47, a potent ‘don’t eat me signal’ is comparatively enriched in EVs upon TH cell activation (Schurch et al., 2017). Phospholipase C gamma 2 (PLCG2), which is associated with inflammatory responses, is highly expressed in rest-EVs (Supplementary Figure 3D) (Tsai et al., 2022; Yu et al., 2005).

Lipidomic analysis was performed to investigate the lipid composition of act-EVs and rest-EVs (Supplementary File 3 & 4). The lipid composition, subcellular origin and physicochemical properties of act-EVs and rest-EVs were largely similar. Lipids like LPE 18:2, LPE 20:1, PC O-30:0, LPE 22:2 and ‘find me signal’ lipid lysophosphatidylcholine such as LPC 40:4-SN2, LPC18:0-SN2, LPC18:0 were differently enriched in act-EVs, whereas PE 38:6, PS 38:4, and PC 12:2_22:1 were differently enriched in rest-EVs (Supplementary Figure 3E-3K).

### TH cells derived EVs trigger upregulation of activation markers on APCs

Supplementary Figure 4A illustrates the gating strategies utilized for the FACS purification of APCs. PBMCs extracted from human blood underwent live/dead staining, and the live cells were utilized for the isolation of B cells and monocytes. T cells were identified through anti-TCR alpha beta and excluded. Monocytes were gated using anti-CD14, while the population negative for both anti-TCR alpha beta and anti-CD14 facilitated the purification of B cells using anti-CD19. To characterize the effect of act-EVs on the activation of APCs, we incubated B-cells and monocytes with different concentrations of EVs in a time series. The optimal concentration of EVs (5x10^10^) needed for the incubation with APCs (2x10^5^ cells seeded at density of 2x10^6^ cells/mL) was determined using titrations, and the optimal time point for incubation was found to be 48 hrs (Supplementary Figure 4B-4D). The LPS levels post EV (Act-EVs and Rest-EVs) isolation from the TH cell culture supernatant were measured and found to be negligible in both (Supplementary Figure 2A).

B cells and monocytes treated with rest-EVs and act-EVs underwent analysis to evaluate the expression status of APC activation markers CD54 (ICAM1), CD40, HLA-DR, and CD83 (HB15) (Figure 3). PMA-Ionomycin (20 ng/ml and 1 μg/ml, respectively) treated cells were used as a positive control and untreated cells were used as a negative control. Both act-EVs and rest-EVs modulates the expression status of activation markers in the recipient APCs. Although, treatment with act-EVs led to a significant increase in CD40 and CD54 (ICAM1) levels were elevated in monocytes. In addition, there was a significant increase in HLA-DR expression in both B cells and monocytes following the treatment with act-EVs compared to rest-EVs. It is intriguing that both rest-EVs and act-EVs induced the upregulation of specific markers, irrespective of the physiological state of the parent TH cells.

**Figure 3:**
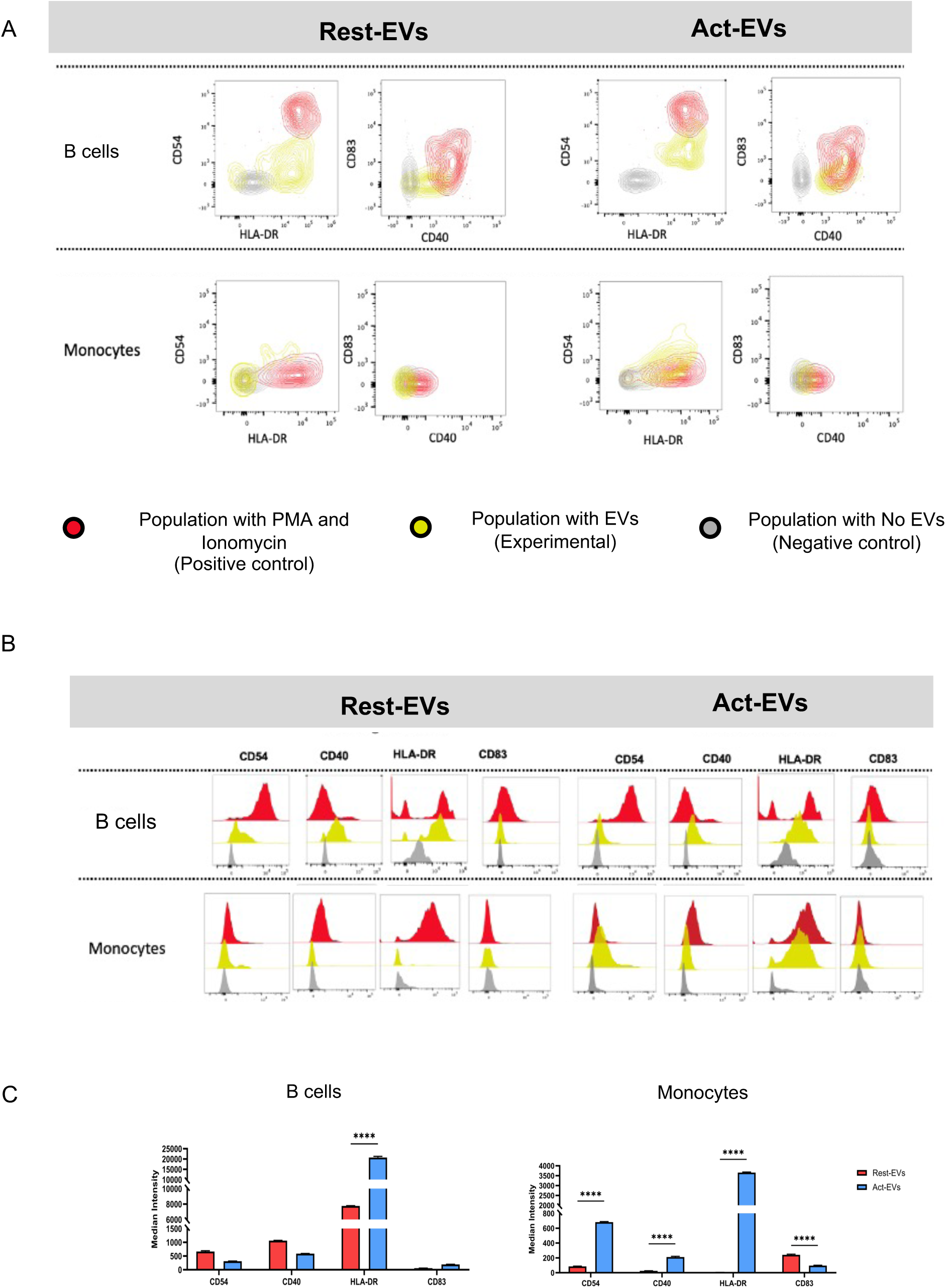
(A) Qualitative analysis of the expression of CD54, CD40, HLA-DR, CD83 expression status by multiparametric cytometer across B cells and monocytes post incubation with rest-EVs and act-EVs. (B) The histogram of positive controls are in red, negative controls are in grey and experimental group are in yellow. (C) The barchart depicts the median intensity of CD54, CD40, HLA-DR, CD83 across B cells and monocytes upon incubation with rest-EVs and act-EVs. Statistical significance was determined using two-way ANOVA Šídák method. The significance level is *p≤0.05, **p≤0.01, ***p≤0.001 and ****p≤0.0001 (n=3 different donors).

### Act-EVs induce a pro-inflammatory cytokine milieu in APCs

To explore the potential of TH cell-derived EVs in triggering pro-inflammatory responses in APCs, these cells were stimulated with both rest-EVs and act-EVs; we then profiled the cytokines and chemokines present in the culture supernatant (Supplementary File 5). Act-EVs selectively induced pro-inflammatory cytokines in APCs, evidenced by the fold change in reference to treatment with rest-EVs (Figure 4A). The secretion of pro-inflammatory cytokines was in concordance with the status of growth factors in B cells and monocytes (Figure 4B). The increased secretion of CCL and CXCL chemokines was primarily observed in B cells following treatment with act-EVs (Figure 4C). Furthermore, interleukins such as IL8, IL5, and IL2 were elevated among APCs after exposure to act-EVs (Figure 4D). These findings underscore that act-EVs were enriched in ligands that induce signalling cascades leading to secretion of pro-inflammatory cytokines and chemokines.

**Figure 4:**
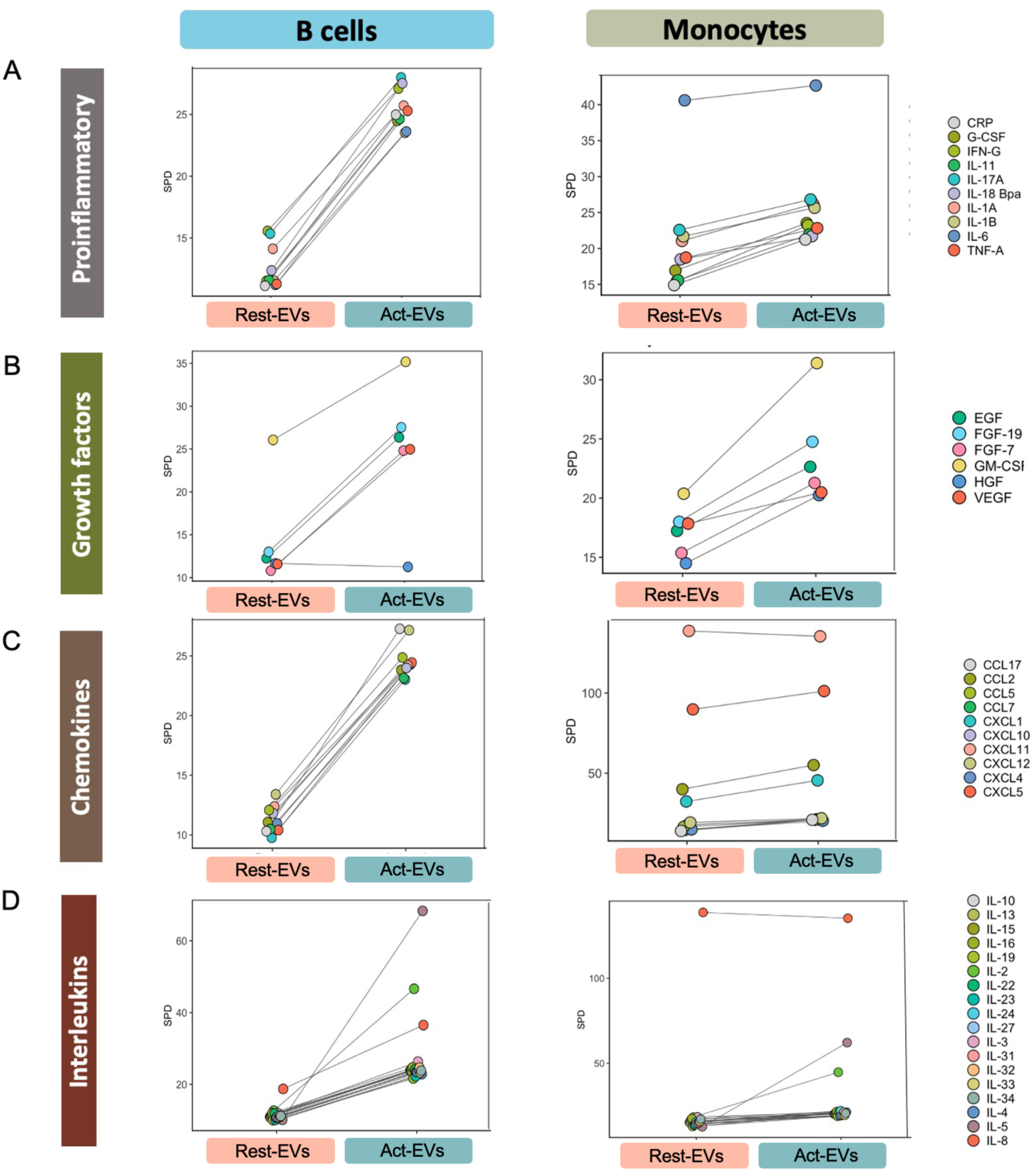
(A-D) Proinflammatory cytokines (A), Growth factors (B), CCL and CXCL chemokines (C), and Interleukins (D) in B cells and monocytes after stimulation with rest-EVs and act-EVs. Data values are plotted as mean values. Cytokines chemokines mean spot densities were calculated by image J (n = 2).

### TH derived EVs reprograms APC in vivo to a pro-inflammatory phenotype

Considering the apparent shift observed in APCs towards a pro-inflammatory state induced by act-EVs in previous in vitro experiments, our subsequent goal was to delve into the repercussions of these act-EVs in vivo. Recognizing that EVs exhibit a rapid pharmacokinetic profile in vivo and can accumulate in the spleen and liver, we hypothesized that the effects of act-EVs would swiftly manifest and be confined to these specific tissues. Furthermore, we expected that the polarization of APCs in the spleen would systematically induce a pro-inflammatory response.

To explore this, FVB mouse splenic TH cells were activated with CD3 beads, and subsequently the EVs were purified 48 hours post-activation (Figure 5A). These purified EVs (act-EVs) were then intravenously injected into FVB mice, and serum and tissue samples were collected 4 hrs post-injection. To account for potential confounding effects resulting from the intervention in animals or the mere injection of EVs, control mice received EVs purified from mouse plasma (plasma-EVs) using a similar procedure. Flow cytometry analysis of the spleen revealed a significant increase in DCs, monocytes, NK cells and neutrophils in mice treated with act-EVs as compared to plasma-EVs (Figure 5C). No significant differences were observed with B cells or T cells -CD4 T cells, CD8 T cells. Importantly, mice injected with act-EVs exhibited a significant increase in GM-CSF, MCP-1, IL-12p70 and IL-1α levels (Figure 5D). This rapid and pronounced surge in serum cytokine levels, coupled with the recruitment of DCs, monocytes, NK cells and neutrophils in the spleen, strongly implies the involvement of activated TH cell exosomes in steering APCs towards a pro-inflammatory phenotype. These findings provide valuable insights into the dynamic interplay between act-EVs and the immune system, particularly in terms of inducing a pro-inflammatory response in vivo.

**Figure 5:**
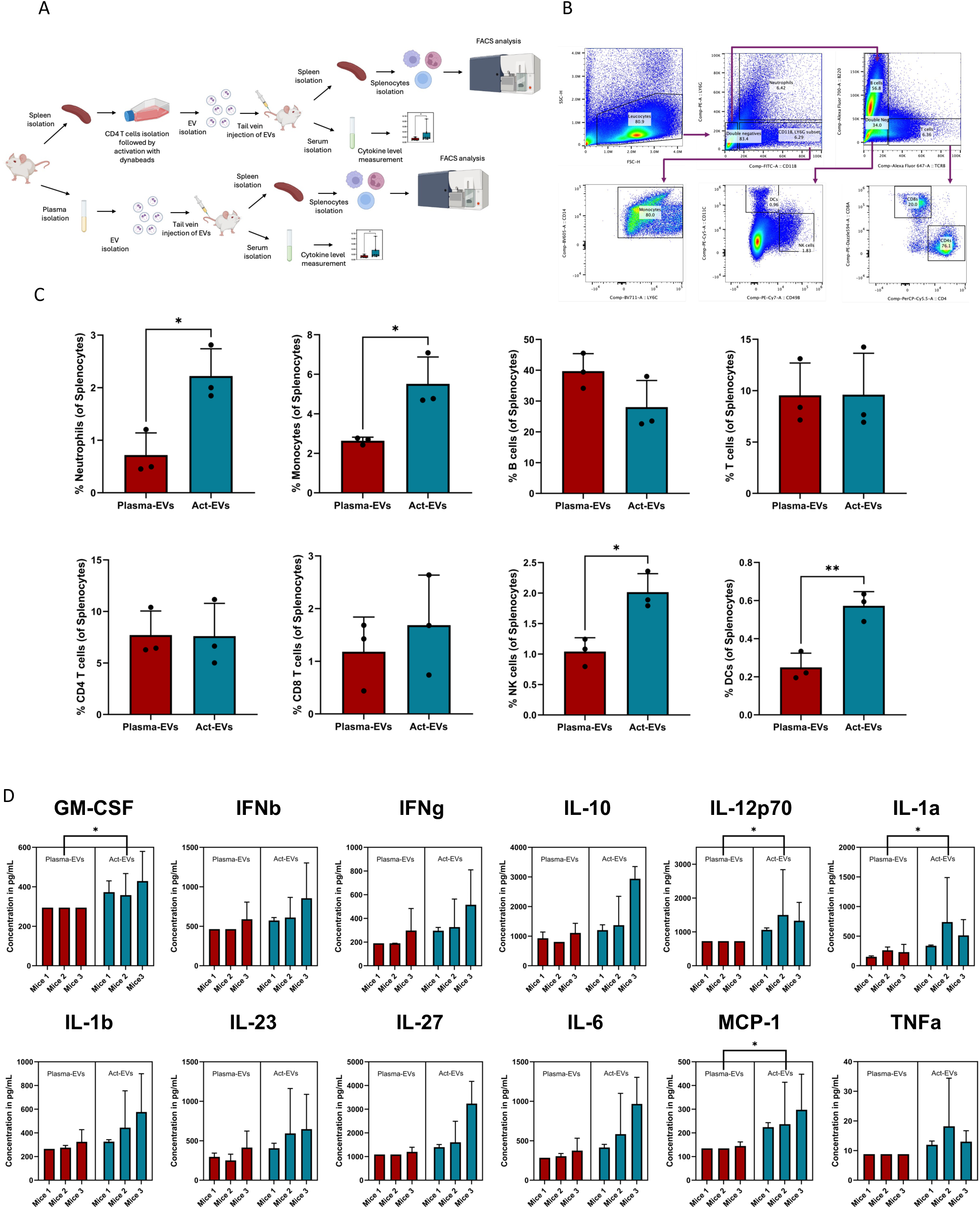
(A) Schematic description of the study aimed at evaluating the physiological response of act-EVs in vivo. (B) Gating strategies of FACS purified of myeloid and lymphocyte cell subsets from mice spleen. (C) Total number of myeloid and lymphocyte cell subsets in spleen from mice (n=3) injected with either act-EVs or plasma-EVs. Unpaired t-test was performed. (D) Serum levels of pro-inflammatory cytokines determined by multiplex cytokine analysis in mice (n= 9, 3 technical replicates for each 3 biological replicates) 4 hours post-injection of either act-EVs or plasma-EVs. Each bar represents a biological replicate, with error bars indicating the standard deviation based on the technical replicates. Nested T test was used to estimate the statistical significance. The significance level is *p≤0.05, **p≤0.01, ***p≤0.001 and ****p≤0.0001.

## Discussion

Recent studies have suggested that TH cell derived EVs may play a role in modulating the immune response during inflammation. Some studies have suggested that TH cell derived EVs may act as decoy vehicles and inhibit the activity of inflammatory cells and cytokines, such as TNF-α and IL-6. Other studies have suggested that TH cell derived EVs may have pro-inflammatory effects, promoting the activation of immune cells and the production of cytokines such as IL-17 (Hazrati et al., 2022; Sun et al., 2021; Sun et al., 2020; Ye et al., 2014). Additionally, it has been reported that TH cell-derived EVs with DGKK trigger apoptosis, oxidative stress, and inflammation through the DAG/PKC/NOX4 pathway (Tu et al., 2023). Overall, the role of TH cell derived EVs in inflammation is still not fully understood and further research is needed to determine their precise mechanisms of action and potential therapeutic applications. This study serves as a resource for understanding the essential mechanism by which TH cell-derived EVs drive pro-inflammatory responses in APCs. By incorporating scRNA seq to study host TH (resting and activated) cells transcriptomics, as well as lipidomics and proteomics to characterize the released EVs and cytokine-chemokine array to study the response of recipient cells upon EV treatment we here report an end-to-end process of cargo transfer. In the context of scRNA sequencing, activated TH cells express high levels of activation markers and effector molecules, and their gene expression profiles are significantly different from those of resting TH cells. TH cells revealed upregulation of transcripts involved in EV biogenesis and the release of a plethora of EVs upon activation.

To test this hypothesis, we purified EVs using SELC and characterised them using NTA analysis, western blotting, nanoFCM, and dSTORM imaging. It was evident the EVs released were heterogenous in both size and cargo contents, and their release increased multifold upon activation. The EVs carried more TCRs and other TH effector molecules like CD69, CD40LG and CD4 upon TH cell activation. The single-particle analysis conducted on EVs showcased clear distinctions in both biochemical and physical properties between rest-EVs and act-EVs. Further, we conducted proteomics of EVs, which revealed enrichment of transmembrane and cytosolic EV marker proteins and TH cell specific proteins. Their content seems to reflect most of the upregulated genes that was expressed in the host TH cells upon activation.

The enrichment of signaling-related proteins, including the calcium-dependent kinase ZAP70, in act-EVs suggests that act-EVs may play a role in cell-cell communication suggesting that act-EVs are primed for the interaction with APCs and modulating T cell responses. Additionally, the enrichment of proteins such as SPN (CD43) and FLNA in act-EVs, further indicates their potential involvement in T cell proliferation, migration, and adhesion dynamics.

The lipidomic analysis provided insights into differences in lipid composition and functional attributes between act-EVs and rest-EVs released by TH cells. While the overall lipid composition was largely similar between act-EVs and rest-EVs, a closer examination revealed nuanced distinctions. Interestingly, lipodomics showed that lysophosphatidylethanolamine, which is generally enriched in activated TH cells, is differentially enriched in act-EVs. Act-EV also had differential expression of lysophosphatidylcholine and apolipoproteins involved in triggering proinflammatory responses (Georgila et al., 2019). Notably, the presence of ’find me signal’ lipids, including certain Lysophosphatidylcholines (LPCs), in act-EVs suggests a potential role in cellular recruitment and communication. Further, enrichment of CD47 (“don’t eat me signal”) seems to ensure their surveillance and reduce the clearance at the site of inflammation (Takimoto et al., 2019).

Further, we wanted to investigate the biochemical message that was being delivered through the EVs from activated TH cells and hence their functional efficacy. Flow cytometric analysis of the EV-treated APC populations showed that EVs isolated from activated TH cells were capable of inducing activation of syngeneic recipient cells. The flow cytometry results demonstrated distinct responses in APCs when exposed to rest-EVs and act-EVs. Act-EVs exhibited a selective induction of pro-inflammatory cytokines in APCs, as evidenced by a substantial fold change in comparison to treatment with rest-EVs. These findings collectively underscore that act-EVs carry a cargo enriched in ligands capable of inducing signaling cascades leading to the secretion of pro-inflammatory cytokines and chemokines. The observed elevation of IL8, IL5, and IL2 highlights the broad pro-inflammatory potential of act-EVs in stimulating various components of the immune response.

Our in vivo investigation into the impact of act-EVs provides compelling evidence of their ability to induce a pronounced pro-inflammatory response. The observed upregulation of pro-inflammatory cytokines, in mice injected with act-EVs, compared to the control group receiving plasma-EVs, underscores the systemic and rapid effects of these vesicles. The preferential accumulation of act-EVs in the spleen and liver aligns with the known tropism of EVs in vivo. The increased presence of DCs, monocytes, NK cells and neutrophils in the spleen following the injection of act-EVs suggests a robust immune response. This heightened activity could be due to various pathways triggered by the cargo molecules within act-EVs or through activation signals to APCs. The possible pathways include CD40-CD40L interactions, RNA-binding proteins, IFN-γ, TNF-α, IL-2, IL-12, and IL-15, all of which are known to promote immune cell recruitment and activation. Although these mechanisms appear to be driving the observed immune response, the exact pathways have not yet been fully elucidated. Similar polarisation in vivo profile is also observed with other immune stimulants such as Lactoferrin (Siqueiros-Cendon et al., 2014). Notably, with the tendency of EVs to enrich in tumours upon systemic administration, act-EVs may play a key role in driving anti-tumour immunity (Gupta et al., 2023). Overall, the observed enrichment of DCs, monocytes, NK cells and neutrophils in the spleen along with elevated levels of GM-CSF, MCP-1, IL-12p70 and IL-1α levels indicates a complex interplay of immunomodulatory signals and antigen-driven responses orchestrated by act-EVs in vivo. This study not only reinforces the role of TH cell-derived EVs in shaping immune responses but also highlights their potential as modulators of inflammation in physiological settings, providing valuable insights for therapeutic interventions targeting immune-related disorders.

In conclusion, we report activated TH EVs can potentially trigger activation of pro-inflammatory response cascades in APCs. Pro-inflammatory responses are a critical element in the progression of several pathologies, including autoimmune diseases. We propose that exploring the EV signalling axis offers a novel approach for characterizing autoimmune diseases, as our findings show EVs deliver cargos to APCs, potentially initiating disease progression. Further, investigating EVs from other T cell subtypes could also improve prognosis in challenging conditions like chronic infections and cancers. The approach described here provides a guideline to perform robust exploratory studies within each cell type and holds a lot of potential in understanding the role of EVs in triggering a plethora of cellular responses.

## Author Contributions

AKJ, MJAW and MLD designed the study. AKJ, NMA, and TT were responsible for analysing scRNA data. GB, SH, and RF worked on LC-MS acquisitions for the lipidomics and proteomics. AKJ and ME performed the proteomics and lipidomics analysis. JC ran the tracking analysis and helped with image analyses. JC and SV helped with Total Internal Reflection Fluorescent Microscopy. ME and RSM worked on flow cytometry data analysis and interpretation. RSM and JC helped with the cytokine array analysis. DG, ME, AL, and SRA carried out the in-vivo study and analysis. AKJ, PFC, MC, LBD, DB, AD, and MLD interpreted results and discussed experiments. AKJ, ME, JC, RSM, and, PFC contributed to the first drafted initial manuscript. MC, LBD, SA, EK, and SV provided insightful feedback and helped in shaping the manuscript. MLD obtained funding for this study. MLD, MJAW and LBD provided infrastructure and supervised the study. All authors contributed to writing and editing the manuscript.

## Acknowledgements

The authors grateful to Dr. Errin Johnson from Sir William Dunn School of Pathology, the University of Oxford, for her help with Electron Microscopy. We are grateful to the MS laboratory at the Target Discovery Institute NDM (Oxford) led by Benedikt M. Kessler for their help and support with Lipidomics and Proteomics. This work was supported by the European commission Horizon 2020 (ERC-2021-SyG_951329 and ERC-2014-AdG_670930), the Kennedy Trust for Rheumatology Research Cell Dynamics Program (KENN202117) and the Wellcome Trust (100262Z/12/Z and 224343/Z/21/Z). Ashwin K. Jainarayanan is supported by the Clarendon Fund, and the Interdisciplinary Bioscience DTP at the University of Oxford (UKRI–BBSRC grant no. BB/M011224/1). Jesusa Capera is a Cancer Research Institute Irvington Fellow supported by the CRI4503 and by a collaborative research grant from Cue Biopharma. Ranjeet Singh Mahla and Pablo F. Céspedes are supported by the Bristol Myers Squibb-Oxford post-doctoral fellowship programme. Pablo F. Céspedes and Michael L. Dustin are supported by Chinese Academy of Medical Sciences Oxford Institute, CAMS Innovation Fund for Medical Sciences (CIFMS) funding code 2018-I2M-2-002.

---

**Supplementary Figure 1:**
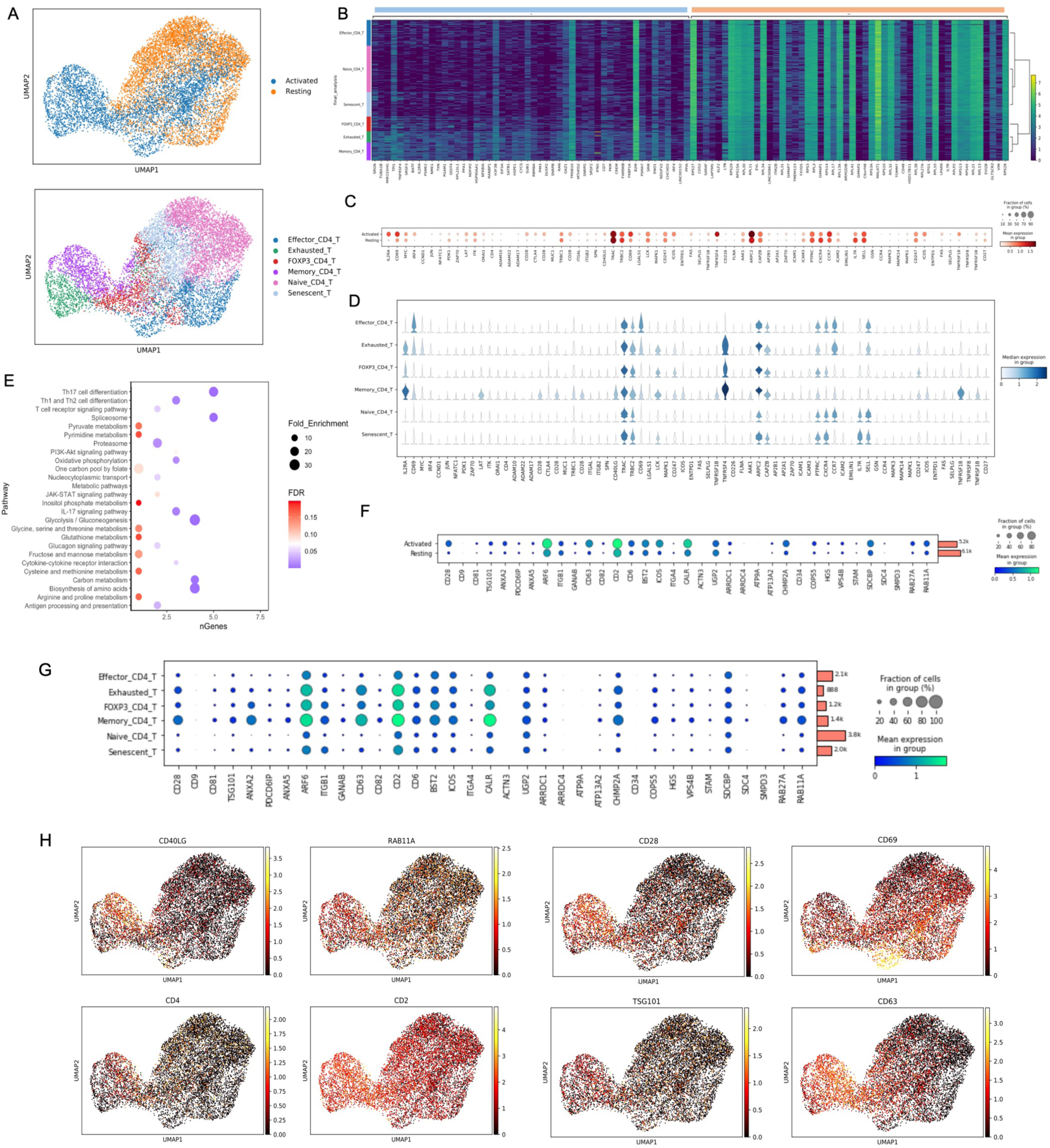
(A) UMAP dimensionality reduction embedding of the activated and resting TH cells (Top panel) and different TH cell types (Bottom panel) coloured by orthogonally generated clusters labelled by manual cell type annotation. (B) Heat map depicting the differentially expressed genes across the different cell types in the activated (Blue) and resting (Orange) TH cell population. (C) Dot plot depicting mean expression (visualized by colour) and fraction of cells expressing (visualized by the size of the dot) key genes involved in T cell activation in resting and activated TH cells. (D) Violin plot of the genes involved in T cell activation across the different TH cell type. (E) Dot plot depicting the KEGG pathways for the top 50 differential expressed genes between the resting and activated TH cells. (F-G) Dot plot depicting mean expression (visualized by colour) and fraction of cells expressing (visualized by the size of the dot) key genes involved in EV biogenesis in resting and activated TH cells (F) and different TH cell type (G). The bars at the right correspond to the total number of cells in that cluster. (H) UMAP dimensionality reduction of the TH cells coloured based on the expression of CD40LG, RAB11A, CD28, CD69, CD4, CD2, TSG101, CD63.

**Supplementary Figure 2:**
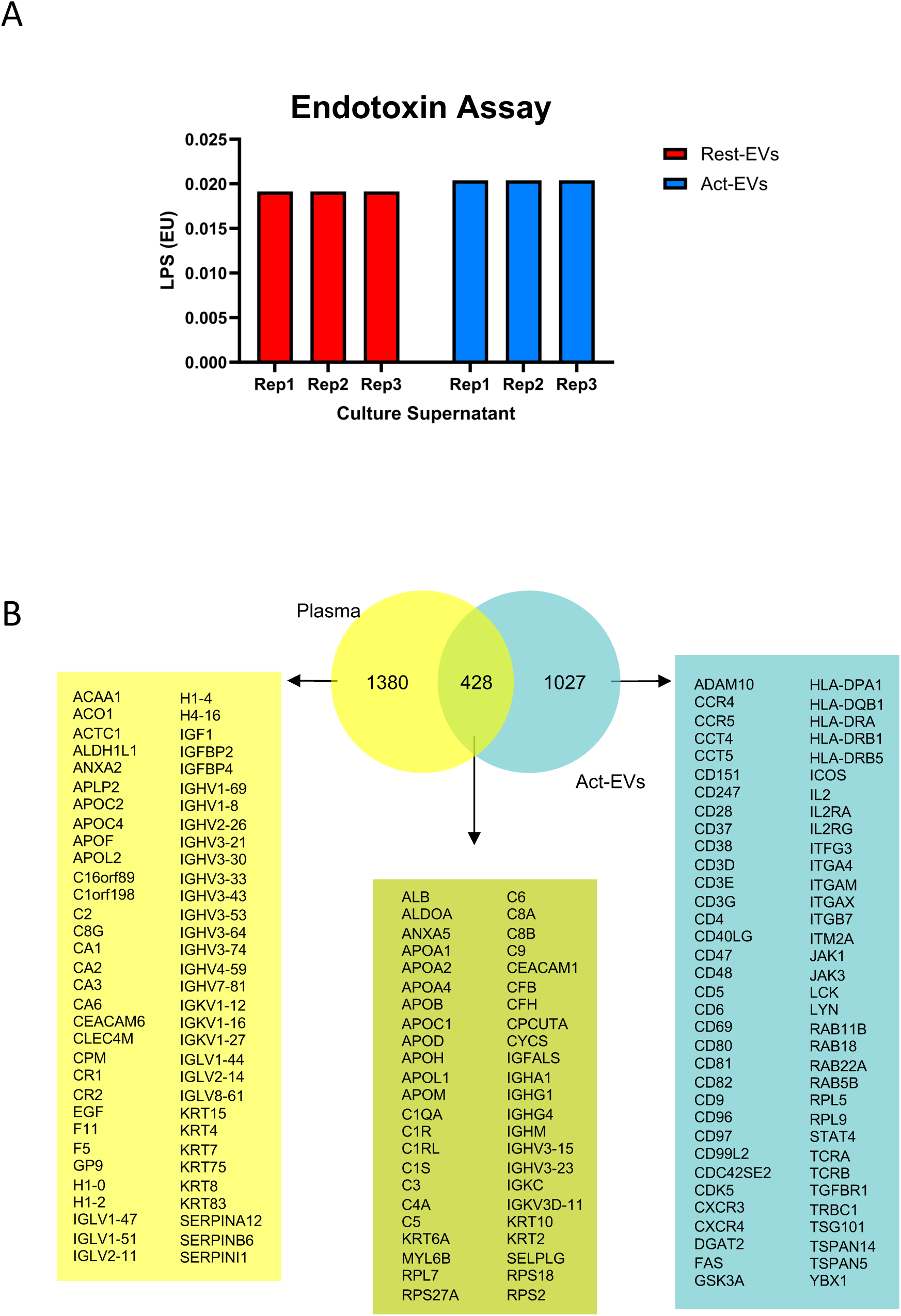
(A) Barchart showing the LPS levels post EV (Act-EVs and Rest-EVs) isolation from the TH cell culture supernatant. (B) Venn diagram depicting the numbers of shared and unique proteins (some of the proteins are listed down) between those detected in act-EVs and enriched in human plasma (J. Proteome Res. 2021, 20, 12, 5241–5263).

**Supplementary Figure 3:**
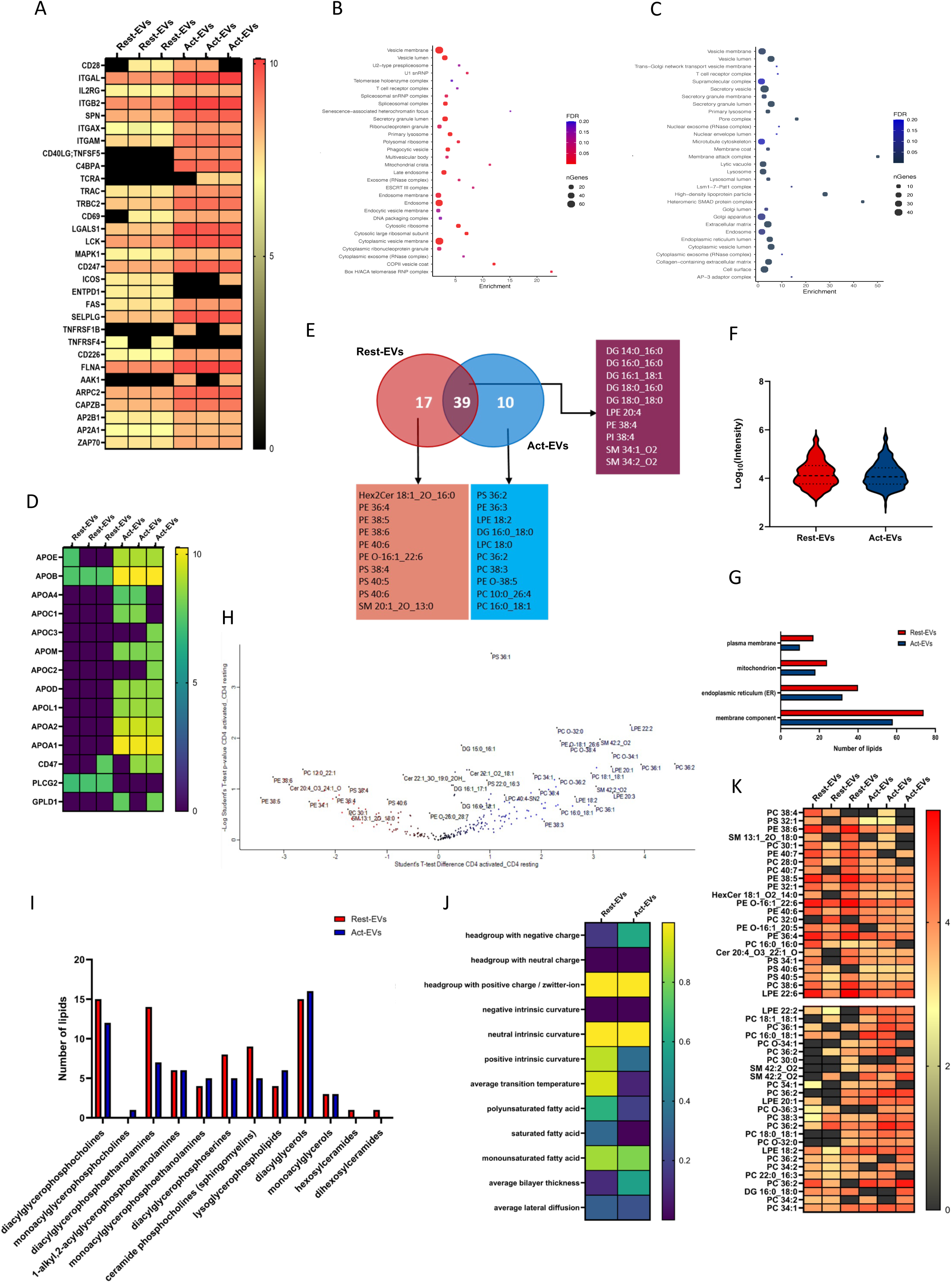
(A) Expression level of signaling related proteins in rest-EVs and act-EVs. (B-C) Gene Ontology (GO) cellular component analysis of act-EVs (B) and rest-EVs (C). (D) Expression level of proteins, including apolipoproteins in rest-EVs and act-EVs. (E) Venn diagram depicting the numbers of shared and unique enriched lipids (some of the lipids are listed down) between the rest-EVs and act-EVs. (F) Violin plot showing the distribution of the lipid contents in the rest-EVs and act-EVs. Lines show the median and quartiles. (G) Bar chart showing the number of enriched lipids -Gene Ontology (GO) cellular component analysis of the enriched lipids found in the rest-EVs and act-EVs. (H) Volcano plot representing the differentially expressed lipids in rest-EVs (Red) and act-EVs (Blue). (I) Barchart depicting the number of lipids under each lipid class found in rest-EVs and act-EVs. (J) Heatmap showing the p-value for different lipid property categories found in the rest-EVs and act-EVs. (K) Heatmap depicting the top 25 differentially enriched lipids from rest-EVs (Top panel) and act-EVs (Bottom panel). LPE - Lysophosphatidylethanolamine, PC - Phosphatidylcholine, PE - Phosphatidylethanolamine, PS - Phosphatidylserine, SM - Sphingomyelin, DG - Diglyceride, LPC - Lysophosphatidylcholine, Cer - Ceramide and PI - Phosphatidylinositol. Data are representative of n =3 individual replicates and the LFQ intensities (for proteins) and intensities (for lipids) were log10 transformed.

**Supplementary Figure 4:**
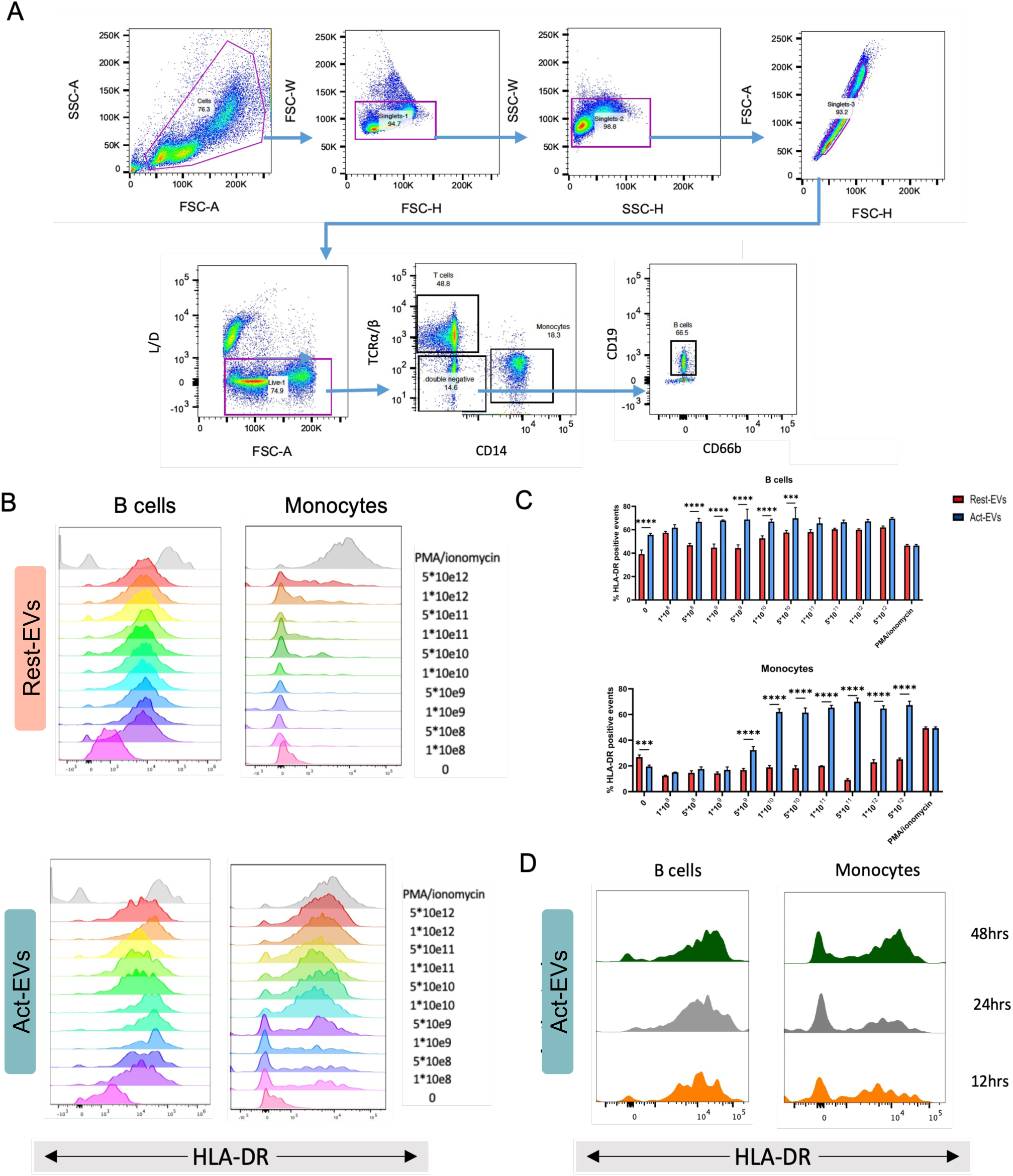
(A) Gating strategies of FACS purified B cells and monocytes from human PBMCs. (B) Titration of rest-EVs and act-EVs concentrations that is required to provoke a response (HLA-DR upregulation) in the donor matched recipient B cells and monocytes. (C) Bar charts showing the percentage of HLA-DR positive events across the B cells and monocytes. Statistical significance was determined using two-way ANOVA Šídák method. (D) Histogram showing incubation duration (12hrs, 24hrs and 48hrs) optimization of act-EVs (5xE10) with donor matched B cells and monocytes. The significance level is *p≤0.05, **p≤0.01, ***p≤0.001 and ****p≤0.0001 (n=3 different donors).

## References

Balint, S., Muller, S., Fischer, R., Kessler, B.M., Harkiolaki, M., Valitutti, S., and Dustin, M.L. (2020). Supramolecular attack particles are autonomous killing entities released from cytotoxic T cells. Science 368, 897–901.

Buzas, E.I. (2023). The roles of extracellular vesicles in the immune system. Nature Reviews Immunology 23, 236–250.

Cai, L., Chao, G., Li, W., Zhu, J., Li, F., Qi, B., Wei, Y., Chen, S., Zhou, G., Lu, X., et al. (2020). Activated CD4(+) T cells-derived exosomal miR-142-3p boosts post-ischemic ventricular remodeling by activating myofibroblast. Aging (Albany NY) 12, 7380–7396.

Cespedes, P.F., Jainarayanan, A., Fernandez-Messina, L., Valvo, S., Saliba, D.G., Kurz, E., Kvalvaag, A., Chen, L., Ganskow, C., Colin-York, H., et al. (2022). T-cell trans-synaptic vesicles are distinct and carry greater effector content than constitutive extracellular vesicles. Nat Commun 13, 3460.

Chen, K., Huang, J., Gong, W., Zhang, L., Yu, P., and Wang, J.M. (2006). CD40/CD40L dyad in the inflammatory and immune responses in the central nervous system. Cell Mol Immunol 3, 163–169.

Colombo, M., Raposo, G., and Thery, C. (2014). Biogenesis, secretion, and intercellular interactions of exosomes and other extracellular vesicles. Annu Rev Cell Dev Biol 30, 255–289.

Deutsch, E.W., Omenn, G.S., Sun, Z., Maes, M., Pernemalm, M., Palaniappan, K.K., Letunica, N., Vandenbrouck, Y., Brun, V., Tao, S.C., et al. (2021). Advances and Utility of the Human Plasma Proteome. J Proteome Res 20, 5241–5263.

Dong, C. (2021). Cytokine Regulation and Function in T Cells. Annu Rev Immunol 39, 51–76

Georgila, K., Vyrla, D., and Drakos, E. (2019). Apolipoprotein A-I (ApoA-I), Immunity, Inflammation and Cancer. Cancers (Basel) 11.

Ginini, L., Billan, S., Fridman, E., and Gil, Z. (2022). Insight into Extracellular Vesicle-Cell Communication: From Cell Recognition to Intracellular Fate. Cells 11.

Gupta, D., Wiklander, O.P.B., Wood, M.J.A., and El-Andaloussi, S. (2023). Biodistribution of therapeutic extracellular vesicles. Extracell Vesicles Circ Nucleic Acids 4, 170–190.

Harris, J.R. (2007). Negative staining of thinly spread biological samples. Methods Mol Biol 369, 107–142.

Hazrati, A., Soudi, S., Malekpour, K., Mahmoudi, M., Rahimi, A., Hashemi, S.M., and Varma, R.S. (2022). Immune cells-derived exosomes function as a double-edged sword: role in disease progression and their therapeutic applications. Biomark Res 10, 30.

Hezel, M.E.V., Nieuwland, R., Bruggen, R.V., and Juffermans, N.P. (2017). The Ability of Extracellular Vesicles to Induce a Pro-Inflammatory Host Response. Int J Mol Sci 18.

Hirsch, Y., Geraghty, J.R., Reiter, C.R., Katz, E.A., Little, C.F., Tobin, M.K., and Testai, F.D. (2023). Unpacking the Role of Extracellular Vesicles in Ischemic and Hemorrhagic Stroke: Pathophysiology and Therapeutic Implications. Transl Stroke Res 14, 146–159.

Hong, X., Schouest, B., and Xu, H. (2017). Effects of exosome on the activation of CD4+ T cells in rhesus macaques: a potential application for HIV latency reactivation. Scientific Reports 7, 15611.

Jainarayanan, A.K., Capera, J., Cespedes, P.F., Conceicao, M., Elanchezhian, M., Thomas, T., Bonner, S., Valvo, S., Kurz, E., Mahla, R.S., et al. (2023). Comparison of different methods for isolating CD8(+) T lymphocyte-derived extracellular vesicles and supramolecular attack particles. J Extracell Biol 2, e74.

Jaiswal, R., and Sedger, L.M. (2019). Intercellular Vesicular Transfer by Exosomes, Microparticles and Oncosomes -Implications for Cancer Biology and Treatments. Front Oncol 9, 125.

Jankovicova, J., Secova, P., Michalkova, K., and Antalikova, J. (2020). Tetraspanins, More than Markers of Extracellular Vesicles in Reproduction. Int J Mol Sci 21.

Jurj, A., Zanoaga, O., Braicu, C., Lazar, V., Tomuleasa, C., Irimie, A., and Berindan-Neagoe, I. (2020). A Comprehensive Picture of Extracellular Vesicles and Their Contents. Molecular Transfer to Cancer Cells. Cancers (Basel) 12.

Kalluri, R. (2024). The biology and function of extracellular vesicles in immune response and immunity. Immunity 57, 1752–1768.

Kalra, H., Drummen, G.P., and Mathivanan, S. (2016). Focus on Extracellular Vesicles: Introducing the Next Small Big Thing. Int J Mol Sci 17, 170.

Kim, K.M., Abdelmohsen, K., Mustapic, M., Kapogiannis, D., and Gorospe, M. (2017). RNA in extracellular vesicles. Wiley Interdiscip Rev RNA 8.

Mkaddem, S.B., Murua, A., Flament, H., Titeca-Beauport, D., Bounaix, C., Danelli, L., Launay, P., Benhamou, M., Blank, U., Daugas, E., et al. (2017). Lyn and Fyn function as molecular switches that control immunoreceptors to direct homeostasis or inflammation. Nat Commun 8, 246.

Mody, P.D., Cannon, J.L., Bandukwala, H.S., Blaine, K.M., Schilling, A.B., Swier, K., and Sperling, A.I. (2007). Signaling through CD43 regulates CD4 T-cell trafficking. Blood 110, 2974–2982.

Muralidharan-Chari, V., Clancy, J., Plou, C., Romao, M., Chavrier, P., Raposo, G., and D’Souza-Schorey, C. (2009). ARF6-regulated shedding of tumor cell-derived plasma membrane microvesicles. Curr Biol 19, 1875–1885.

Perez-Villar, J.J., Whitney, G.S., Bowen, M.A., Hewgill, D.H., Aruffo, A.A., and Kanner, S.B. (1999). CD5 negatively regulates the T-cell antigen receptor signal transduction pathway: involvement of SH2-containing phosphotyrosine phosphatase SHP-1. Mol Cell Biol 19, 2903–2912.

Ruiz, M., Frej, C., Holmer, A., Guo, L.J., Tran, S., and Dahlback, B. (2017). High-Density Lipoprotein-Associated Apolipoprotein M Limits Endothelial Inflammation by Delivering Sphingosine-1-Phosphate to the Sphingosine-1-Phosphate Receptor 1. Arterioscler Thromb Vasc Biol 37, 118–129.

Saliba, D.G., Cespedes-Donoso, P.F., Balint, S., Compeer, E.B., Korobchevskaya, K., Valvo, S., Mayya, V., Kvalvaag, A., Peng, Y., Dong, T., et al. (2019). Composition and structure of synaptic ectosomes exporting antigen receptor linked to functional CD40 ligand from helper T cells. Elife 8.

Savinko, T., Guenther, C., Uotila, L.M., Llort Asens, M., Yao, S., Tojkander, S., and Fagerholm, S.C. (2018). Filamin A Is Required for Optimal T Cell Integrin-Mediated Force Transmission, Flow Adhesion, and T Cell Trafficking. J Immunol 200, 3109–3116.

Scanu, A., Molnarfi, N., Brandt, K.J., Gruaz, L., Dayer, J.-M., and Burger, D. (2008). Stimulated T cells generate microparticles, which mimic cellular contact activation of human monocytes: differential regulation of pro-and anti-inflammatory cytokine production by high-density lipoproteins. Journal of Leukocyte Biology 83, 921–927.

Schneider, C.A., Rasband, W.S., and Eliceiri, K.W. (2012). NIH Image to ImageJ: 25 years of image analysis. Nature Methods 9, 671–675.

Schurch, C.M., Forster, S., Bruhl, F., Yang, S.H., Felley-Bosco, E., and Hewer, E. (2017). The "don’t eat me" signal CD47 is a novel diagnostic biomarker and potential therapeutic target for diffuse malignant mesothelioma. Oncoimmunology 7, e1373235.

Sheehan, C., and D’Souza-Schorey, C. (2019). Tumor-derived extracellular vesicles: molecular parcels that enable regulation of the immune response in cancer. J Cell Sci 132.

Siqueiros-Cendon, T., Arevalo-Gallegos, S., Iglesias-Figueroa, B.F., Garcia-Montoya, I.A., Salazar-Martinez, J., and Rascon-Cruz, Q. (2014). Immunomodulatory effects of lactoferrin. Acta Pharmacol Sin 35, 557–566.

Sun, J., Jia, H., Bao, X., Wu, Y., Zhu, T., Li, R., and Zhao, H. (2021). Tumor exosome promotes Th17 cell differentiation by transmitting the lncRNA CRNDE-h in colorectal cancer. Cell Death Dis 12, 123.

Sun, P., Wang, N., Zhao, P., Wang, C., Li, H., Chen, Q., Mang, G., Wang, W., Fang, S., Du, G., et al. (2020). Circulating Exosomes Control CD4(+) T Cell Immunometabolic Functions via the Transfer of miR-142 as a Novel Mediator in Myocarditis. Mol Ther 28, 2605–2620.

Szabo, P.A., Levitin, H.M., Miron, M., Snyder, M.E., Senda, T., Yuan, J., Cheng, Y.L., Bush, E.C., Dogra, P., Thapa, P., et al. (2019). Single-cell transcriptomics of human T cells reveals tissue and activation signatures in health and disease. Nat Commun 10, 4706.

Takimoto, C.H., Chao, M.P., Gibbs, C., McCamish, M.A., Liu, J., Chen, J.Y., Majeti, R., and Weissman, I.L. (2019). The Macrophage ‘Do not eat me’ signal, CD47, is a clinically validated cancer immunotherapy target. Annals of Oncology 30, 486–489.

Tenger, C., and Zhou, X. (2003). Apolipoprotein E modulates immune activation by acting on the antigen-presenting cell. Immunology 109, 392–397.

Thery, C., Witwer, K.W., Aikawa, E., Alcaraz, M.J., Anderson, J.D., Andriantsitohaina, R., Antoniou, A., Arab, T., Archer, F., Atkin-Smith, G.K., et al. (2018). Minimal information for studies of extracellular vesicles 2018 (MISEV2018): a position statement of the International Society for Extracellular Vesicles and update of the MISEV2014 guidelines. J Extracell Vesicles 7, 1535750.

Tinevez, J.Y., Perry, N., Schindelin, J., Hoopes, G.M., Reynolds, G.D., Laplantine, E., Bednarek, S.Y., Shorte, S.L., and Eliceiri, K.W. (2017). TrackMate: An open and extensible platform for single-particle tracking. Methods 115, 80–90.

Torregrosa Paredes, P., Esser, J., Admyre, C., Nord, M., Rahman, Q.K., Lukic, A., Radmark, O., Gronneberg, R., Grunewald, J., Eklund, A., et al. (2012). Bronchoalveolar lavage fluid exosomes contribute to cytokine and leukotriene production in allergic asthma. Allergy 67, 911–919.

Tsai, A.P., Dong, C., Lin, P.B.-C., Messenger, E.J., Casali, B.T., Moutinho, M., Liu, Y., Oblak, A.L., Lamb, B.T., Landreth, G.E., et al. (2022). PLCG2 is associated with the inflammatory response and is induced by amyloid plaques in Alzheimer’s disease. Genome Medicine 14, 17.

Tu, G.W., Zhang, Y., Ma, J.F., Hou, J.Y., Hao, G.W., Su, Y., Luo, J.C., Sheng, L., and Luo, Z. (2023). Extracellular vesicles derived from CD4(+) T cells carry DGKK to promote sepsis-induced lung injury by regulating oxidative stress and inflammation. Cell Mol Biol Lett 28, 24.

van der Vlist, E.J., Arkesteijn, G.J., van de Lest, C.H., Stoorvogel, W., Nolte-’t Hoen, E.N., and Wauben, M.H. (2012). CD4(+) T cell activation promotes the differential release of distinct populations of nanosized vesicles. J Extracell Vesicles 1.

Willms, E., Cabanas, C., Mager, I., Wood, M.J.A., and Vader, P. (2018). Extracellular Vesicle Heterogeneity: Subpopulations, Isolation Techniques, and Diverse Functions in Cancer Progression. Front Immunol 9, 738.

Ye, S.B., Li, Z.L., Luo, D.H., Huang, B.J., Chen, Y.S., Zhang, X.S., Cui, J., Zeng, Y.X., and Li, J. (2014). Tumor-derived exosomes promote tumor progression and T-cell dysfunction through the regulation of enriched exosomal microRNAs in human nasopharyngeal carcinoma. Oncotarget 5, 5439–5452.

Yu, P., Constien, R., Dear, N., Katan, M., Hanke, P., Bunney, T.D., Kunder, S., Quintanilla-Martinez, L., Huffstadt, U., Schroder, A., et al. (2005). Autoimmunity and inflammation due to a gain-of-function mutation in phospholipase C gamma 2 that specifically increases external Ca2+ entry. Immunity 22, 451–465.

Zewinger, S., Reiser, J., Jankowski, V., Alansary, D., Hahm, E., Triem, S., Klug, M., Schunk, S.J., Schmit, D., Kramann, R., et al. (2020). Apolipoprotein C3 induces inflammation and organ damage by alternative inflammasome activation. Nature Immunology 21, 30–41.

Zhang, H., Freitas, D., Kim, H.S., Fabijanic, K., Li, Z., Chen, H., Mark, M.T., Molina, H., Martin, A.B., Bojmar, L., et al. (2018). Identification of distinct nanoparticles and subsets of extracellular vesicles by asymmetric flow field-flow fractionation. Nat Cell Biol 20, 332–343.

Zirlik, A., Maier, C., Gerdes, N., MacFarlane, L., Soosairajah, J., Bavendiek, U., Ahrens, I., Ernst, S., Bassler, N., Missiou, A., et al. (2007). CD40 ligand mediates inflammation independently of CD40 by interaction with Mac-1. Circulation 115, 1571–1580.

